# An effective and safe maize seed chipping protocol using clipping pliers with applications in small-scale genotyping and marker-assisted breeding

**DOI:** 10.1101/2024.04.01.587552

**Authors:** Brian Zebosi, John Ssengo, Lander F Geadelmann, Erica Unger-Wallace, Erik Vollbrecht

**Affiliations:** Department of Genetics, Development and Cell Biology, Iowa State University, Ames, Iowa 50011, USA; Genetics and Genomics Graduate Program, Iowa State University, Ames, Iowa 50011, USA; Department of Plant Pathology, Entomology and Microbiology, Iowa State University, Ames, Iowa, 50011 USA; Plant Biology Graduate Program, Iowa State University, Ames, Iowa 50011, USA

**Keywords:** Maize, Genotyping, DNA extraction, Seed chipping

## Abstract

In applications such as marker-assisted breeding and positional cloning, tissue sampling and plant tracking are vital steps in the genotyping pipeline. They enable the identification of desirable seedlings, saving time and reducing the cost, space, and handling required for growing adult plants, especially for greenhouses and winter nurseries. Small-scale marker-assisted selection laboratories rely heavily on leaf-based genotyping, which involves over-planting large, segregating populations followed by leaf sampling, genotyping, and backtracking to identify desired individuals, which is costly and laborious. Thus, there is a need to adopt seed-based genotyping to reduce costs and save time. Therefore, we developed a safe and cheap seed-chipping protocol using clipping pliers to chip seeds to genotype before planting. To identify a cost-effective and high throughput DNA extraction method, we tested four extraction methods and assessed the quality of the seed DNA using PCR. For three of the methods, seed-based DNA was of comparable quality to DNA extracted from leaf punches. We also compared seed- and leaf-derived DNA from the same individuals in a segregating population to test for genotyping miscalls that could arise due to the presence of maternally derived pericarp in the seed samples and found zero miscalls of 43 potential instances. Germination rates of chipped and unchipped seeds were the same for the inbreds tested, B73 and Mo17. However, chipped seeds grew slower until ∼14 days after sowing. Overall, seed sampling using clipping pliers provides a simple, reliable, and high-throughput method to identify specific genotypes before planting.

**Key Features:**

- Provides a quick, safe, and cheap sampling technique for maize kernels that may also be suitable for other plants with relatively large seeds.
- Includes procedures and materials to track and organize samples within and across batches involving hundreds to thousands of seeds.
- Seeds can be sampled and genotyped relatively quickly for planting; in one day 384 seeds can be sampled, processed for DNA, and genotyped by PCR.

## [Background]

Over the last decades, marker-assisted selection (MAS) has become integral to most plant breeding systems because quantitative trait loci (QTL) and molecular markers linked to desired traits have enhanced the selection efficiency of desirable plants and shortened breeding cycles (Xu & Crouch, 2008). Unlike commercial breeding companies with highly automated and streamlined MAS and trait discovery programs, public-sector breeding programs and small-scale genotyping laboratories rely on laborious and time-consuming tissue sampling and DNA extraction procedures (Gao et al., 2008). Compared to the proprietary seed-based MAS programs utilized by industry scientists, public sector genotyping systems rely heavily on leaf-based DNA genotyping, which is resource- and time-demanding (Xu & Crouch, 2008). Thus, to speed up MAS and related approaches, there is a need to adopt seed-based genotyping, which reduces labor costs and saves space and time through selection of desirable genotypes during the off-season (Gao et al., 2008; Xu & Crouch, 2008). During trait discovery, seed-based DNA genotyping ensures that seeds with desirable traits are selected and planted, thus saving time and costs associated with greenhouse use and winter fields. However, in addition to tissue sampling, optimizing rapid and cost-effective DNA extraction methods to screen large populations will be essential for a successful MAS and genotyping pipeline.

Seed-based DNA genotyping methods have been implemented in small-scale maize breeding programs and genotyping laboratories but are still ineffective based on safety and scalability (Gao et al., 2008; Mills et al., 2020). One seed-based DNA genotyping method is based on soaking maize kernels in water before endosperm chipping (Gao et al., 2008). The caveats of this procedure include susceptibility to fungal contamination and low seed germination if kernels are not dried and stored well. Another common method uses a razor blade for seed chipping (Mills et al., 2020), which presents injury hazards and is time-consuming, thus making it impractical to chip a large population quickly. In this study, we used clipping pliers, a safe alternative to previous chipping methods that is scalable to high throughput applications. This seed chipping method did not significantly impact germination rates for B73 and Mo17 seeds, although seedlings from chipped seeds had reduced vigor that was evident for 14 days after planting, after which time they appeared comparable to their non-chipped sibling seedlings. In addition, we tested four DNA extraction methods, one each based on: cetyltrimethylammonium bromide (CTAB) (Yi et al., 2018), sodium dodecyl sulfate (SDS) (Edwards et al., 1991), urea (Leach et al., 2016), and guanidine hydrochloride (Gao et al., 2010), and optimized a cost- and time-efficient DNA extraction protocol for maize endosperm samples.

### Materials and Reagents

#### Biological materials

1. Maize kernels (seeds)

### Laboratory Supplies

1. 2.2 mL 96-well sample collection plate (VWR, catalog number 43001-0020)
2. 1.2 mL sampling tubes (VWR, catalog number 83009-678)
3. 2.0 mL safe-lock microcentrifuge tubes (Eppendorf, catalog number 022363352)
4. Labeling tape

### Equipment

1. Forceps (VWR, catalog number 82027-446)
2. Pet nail clippers (Resco, part number [PF0728](https://www.rescopetproducts.com/product-page/resco-original-guillotine-nail-trimmer-small-medium-1); see General Note 1)
3. Mini ice cube tray, used for seed storage, 135 cubes 0.5 inch size (e.g. [Source 1](https://www.walmart.com/ip/Mini-Ice-Cube-Tray-with-Lid-and-Bin-TINANA-135-4-PCS-Small-Ice-Trays-for-Freezer-Easy-Release-Nugget-Ice-Tray-with-Lid-Green/5168670900) or [Source 2](https://www.amazon.com/FDDBI-Mini-Cube-Tray-Freezer/dp/B0C5T34JGC)).

### Software

1. The Tissue Sample Plate Mapper software is available as a Google sheet plug-in (Google Workspace Marketplace, plug-in number 115639436496).
2. Excel or other spreadsheets may be useful for additional notetaking.
3. Primer3 software (https://bioinfo.ut.ee/primer3-0.4.0/)
4. MaizeGDB (http://www.maizegdb.org) for primer blast analysis

### Procedure

A. **Labeling seed trays and collection plates**
  1. During seed chipping, two 96-well sample collection plates are needed. One plate (the tube holder) holds one sampling tube at a time to aid tissue collection during chipping.
  2. The second plate (for tissue storage) stores up to 96 accumulated sampling tubes with chipped tissue. Optionally, prepare the tissue storage plate by printing a plate map using the tissue-sample plate mapper software, overlaying the map on a 96-well sample collection plate and securing it with clear tape (see General Note 4). Label this plate with a unique ID using tape and a marker.
  3. One seed storage tray is used to hold chipped maize seeds until analysis and decision-making are completed. Label this tray with the same unique ID to pair it with a tissue storage plate.
B. **Maize seed chipping** See Video 1 for a demonstration of the complete chipping procedure.
  1. On a large laboratory bench, place the tube-holder plate (Figure 1C), labeled tissue storage plate (Figure 1B), and seed storage tray (Figure 1A) with the tube-holder plate closest to you and the seed storage tray farthest from you.
  2. Place one sampling tube in one of the tube holder plate’s bottommost corner wells. The corner location facilitates positioning the chipper right above the sampling tube (Figure 2H, 2K).
  3. Place one sampling tube in one of the tube holder plate’s bottommost corner wells. The corner location facilitates positioning the chipper right above the sampling tube (Figure 2H, 2K).
  4. Using the dominant hand hold the chipper blade side u, with the thumb on the top handle and the other fingers holding the lower handle with the spring attached that moves the blade.
  5. With your other hand’s thumb and index finger, pick up and hold one maize seed by the tip cap with the embryo side facing toward the thumb (Figure 2H-I).
  6. While holding the chipper orifice directly and slightly above the sampling tube in the tube holder plate, hold the seed approximately perpendicular to the pliers and place its crown (Figure 2A) into the chipper hole and gently chip the seed repeatedly to shave off fine tissue particles (Figures 2H-J). For example, three to five passes of the chipping blade are usually required with B73 and Mo17 kernels. **Critical:** Chip and shave the tissue finely (Figures 2B, 2D) because large chip particles (Figure 2F) grind poorly and barely yield extracted DNA (see General Note 2). **Critical:** It is imperative to avoid damaging the embryo (Figure 2A) while chipping the endosperm. Secondary chips may be collected if necessary, for example from small or rounded kernels like from the W22 inbred, by slightly tilting the seed to chip and shave fine endosperm particles as described above, but this time at a 45 degree angle to the kernel axis (Figure 2K-M). Combined primary and secondary chipping also increases the proportion of endosperm tissue in the sample relative to maternal pericarp tissue. **Critical:** Processing too much tissue may reduce DNA yield. Approximately 15-30 mg of tissue is sufficient to yield DNA for use as PCR templates using the extraction methods tested.
  7. If the SDS, filter plate or Urea method is used for DNA extraction (Part C) then remove the sample tube with chipped tissue from the tube-holder plate and place it into the tissue storage plate; then place the chipped seed into the appropriate well in the seed tray. For tracking purposes, match the chipped seed’s position in the seed tray grid with the position of the sampling tube on the tissue storage plate grid. If CTAB is used for DNA extraction (Part C) then transfer the chipped tissue into a labeled 2 mL microcentrifuge tube instead.
  8. Clean the clipping pliers by wiping with a dry tissue and/or blowing off any chip residue.
  9. Repeat steps two to eight with a new sampling tube and kernel until 96 or the desired number of samples have been collected.
  10. Proceed to DNA extraction.
C. **DNA extraction options**
  1. Guanidine-HCl filter plate extraction protocol (see Supplemental protocol 1)
  2. SDS-Based DNA Extraction Protocol (see Supplemental protocol 2)
  3. Urea-Based DNA Extraction Protocol (see Supplemental protocol 3)
  4. Modified CTAB DNA Extraction protocol with steel-bead (Yi et al., 2018).
D. **PCR and Genotyping**
  1. Design high-quality genotyping markers using the maize reference genome (http://www.maizegdb.org) and Primer3 (https://bioinfo.ut.ee/primer3-0.4.0/).
  2. Set up a 13 μl PCR reaction composed of 2X GoTaq GreenMaster Mix (6.5 μl), DMSO (0.5 μl), forward and reverse primer mix (1 μl of 5 μM each), water (2 μl), and DNA (3 μl). For amplifying GC-rich regions, substitute the water in the PCR reaction with betaine (2.5 μl of 5M).
  3. Run standard PCR cycling parameters according to the primers used and product size. For instance, for Figure 4 the primers used were BZ316 (ATCTGCATCCTGCGACGCAAC) and BZ317 (GTCGGCGGTCTTTCTCGAGTC) and PCR conditions were 36 cycles of 94 °C for 30 seconds, 60°C for 30 seconds and 72 °C for 60 seconds. This reaction produced amplicon sizes of 555 bp (B73) and 416 bp (Mo17).
  4. Run the PCR-amplified products on agarose gels for genotyping.
  5. Record the genotypes. If using the tissue-sample plate mapper software then genotype calls may be entered in the ordered Sample List tab generated by the software.
  6. After data analysis, use forceps to carefully pick chipped seeds with the genotype(s) of interest from the seed storage trays.

**Figure 1.**
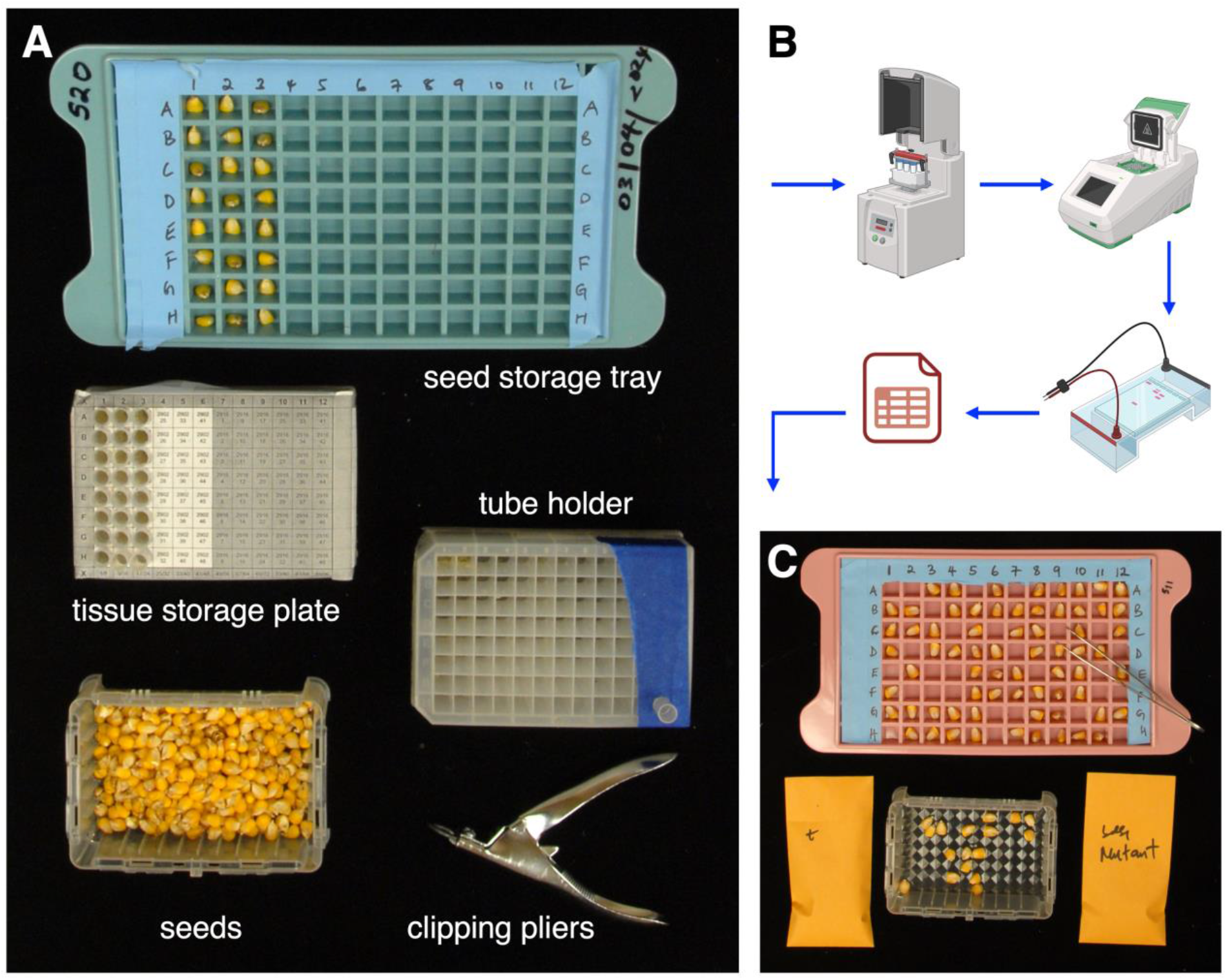
Maize seed chipping setup and genotyping. (A) Sampling includes nail clipping pliers, seeds, a 96-well plate acting as a tube holder plate, a 96-well tissue storage plate overlaid with a sampling map to accumulate chipped samples in tubes, and a seed storage tray. (B) Once chipping is completed, chipped tissue samples are processed for DNA extraction which is genotyped using PCR and an agarose gel and data analysis. (C) With data tracking, seeds with genotype of interest are selected for downstream applications.

**Figure 2.**
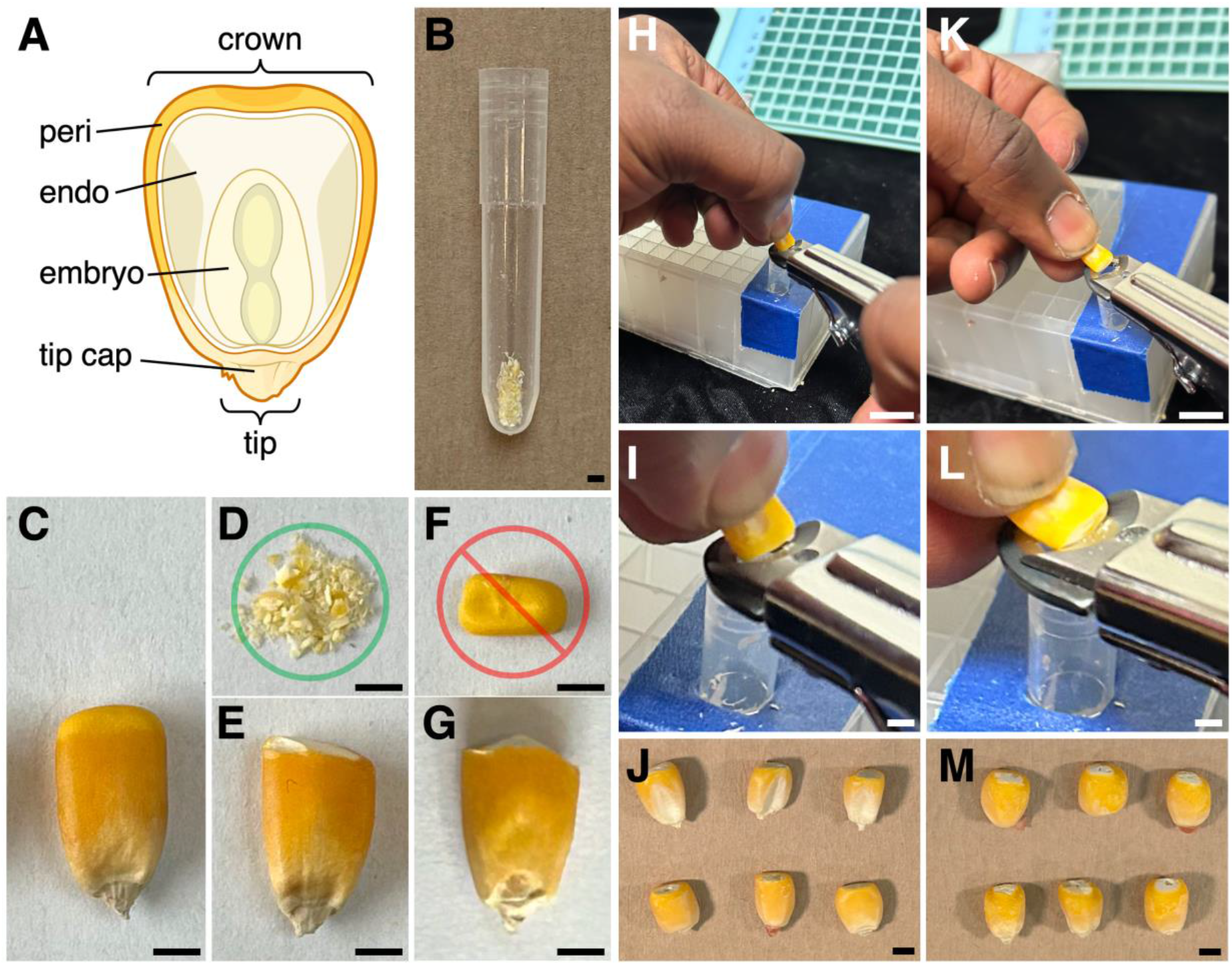
Maize kernels and the seed chipping process. (A) Anatomy of a maize kernel in longitudinal section; peri, pericarp; endo, endosperm. (B) An appropriate volume of seed chips in a collection tube. (C) Intact, unchipped kernel (D-E) properly chipped kernel shavings (D, green circle) removed from the crown of the kernel (E). (F-G) Improperly chopped, large chunk(s) (F, red circle) removed from the crown (G). (H-J) Seed chipping of the crown is done with the embryo facing you and the seed nearly perpendicular to the clipping pliers. (K-M) Secondary chips, e.g. from round-shaped kernels, produced by holding the seed with embryo facing upwards and nearly parallel to the chipping pliers’ blade, to shave chips from the back of the seed, i.e., at a 45-degree angle to the first chipping surface. Scale bars 2 mm except (H, K) scale bars 10 mm.

### Data analysis and validation of protocol

Seed chipping with nail clipping pliers provides safer, scalability and better throughput than razor blade-based methods. 30-40 minutes are required to chip 96 maize seeds, depending on seed size and shape. Smaller or round seeds are challenging to hold and need more time to chip than larger ones. To examine the effects of seed chipping on germination, we planted chipped and unchipped seeds from the same seed packet in three replications of 32 in the greenhouse and scored germination and vigor. We noticed that chipping did not impact the germination rate, but chipped seeds initially grew slower after planting (Figure 3; see General Note 3).

**Figure 3.**
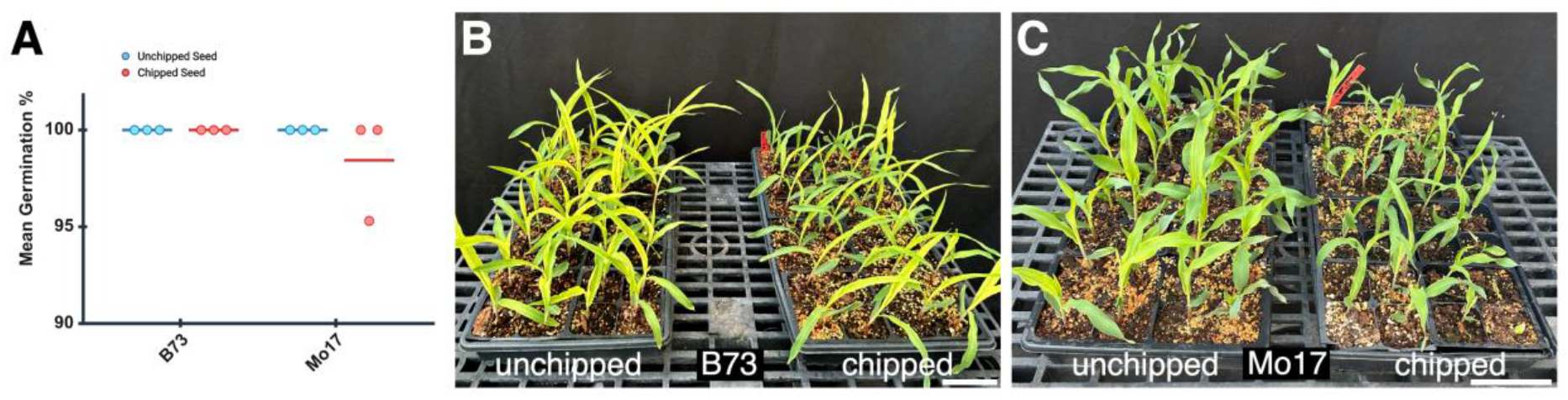
Effect of seed chipping on germination and seedling growth across B73 and Mo17 maize inbreds. (A) Germination frequency of unchipped and chipped seeds across two maize inbreds, Mo17 and B73. Each dot indicates the percentage germination from one replicate of 32 sown kernels. (B) Post-germination seedlings at 14 days to compare growth of unchipped (left) and chipped seeds (right) for B73. (C) As in (B) but for Mo17. Scale bars 10 cm.

We evaluated quality of DNA extracted from seeds by comparing it to leaf-extracted DNA in a PCR test across the different DNA extraction methods. The results indicated that the quality of the DNA extracted from seeds and leaves was comparable using three of the methods tested (Figure 4A). The exception was the urea-based method, where DNA extracted from seed chips performed poorly (Figure 4A). Regarding usability and high throughput, the extraction methods are user-friendly and high throughput except for CTAB. For instance, with CTAB extraction we process a maximum of 48 samples at a time, whereas with other extraction methods 384 samples can be processed at a time once chipping is complete. The guanidine-HCl with GF/F filter plate method has produced the best quality DNA but can include significant expense for reagents and the filter plate. Thus, in our tests the SDS extraction method optimized the combination of usability, high throughput, and cost-effectiveness (e.g. figure 4B).

**Figure 4.**
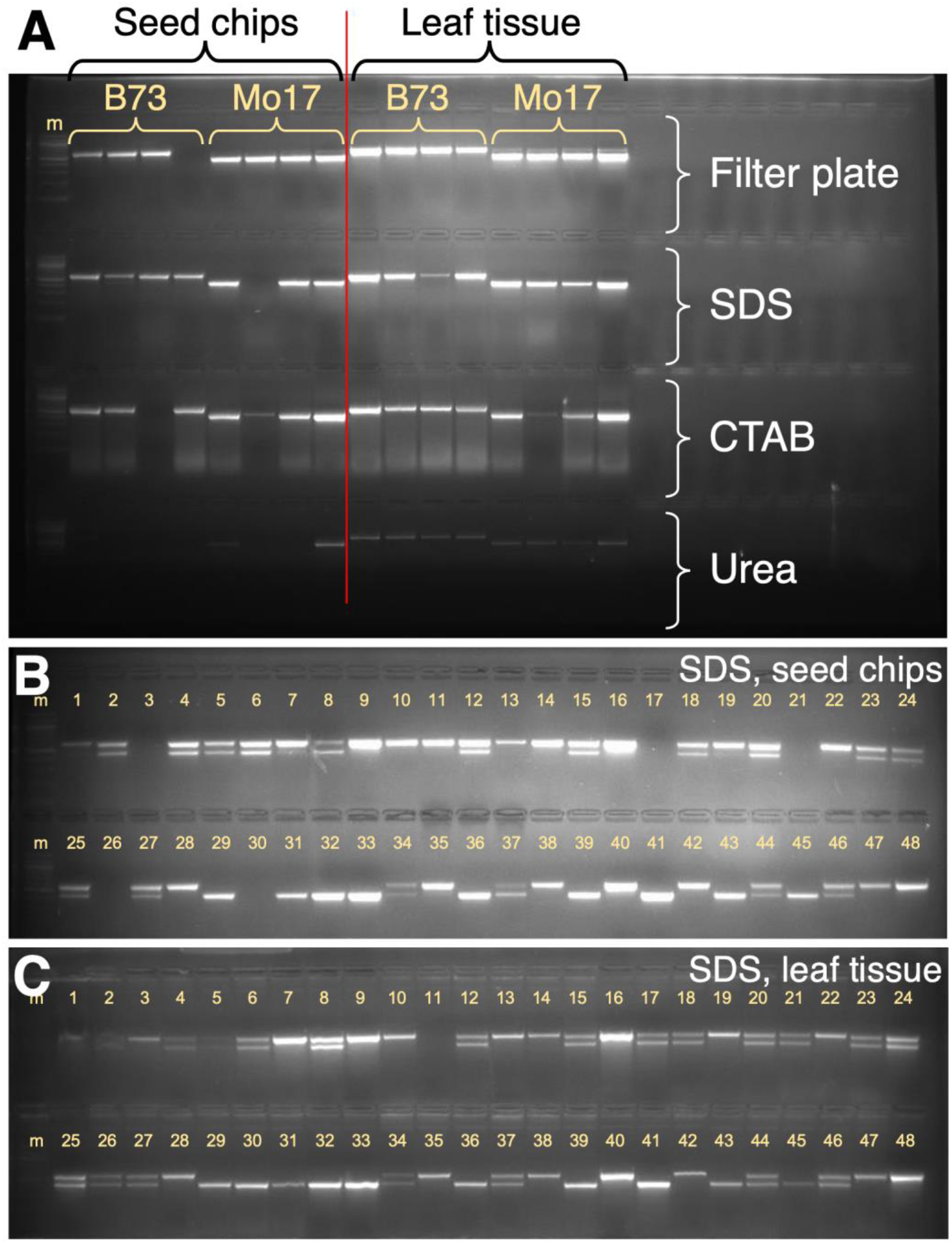
PCR-based genotyping using DNA extracted from chipped seed endosperm, and comparison to DNA from leaves. (A) Comparison of seed-based and leaf-based DNA across four DNA extraction methods. (B-C) Testing for pericarp-based genotyping miscalls by comparing corresponding chipped seed-(B) and leaf-(C) based DNA from the same 96 individuals; representative data from 48 individuals (1-48, yellow numbers) are shown. Homozygotes show upper band alone (e.g. individual 7) or lower band alone (e.g. individual 29); heterozygotes show both bands (e.g. individual 6). m, marker DNA ladder.

In maize seed-based genotyping, maternal pericarp tissue has led to incorrect genotype results. Gao et al. (2008) reported that among the homozygotes within an F2 population, false heterozygous genotyping errors occurred at a rate of 3.8% due to maternal pericarp DNA when chipping was performed after soaking maize kernels in water. To minimize pericarp contribution, we included secondary chips which enriches chip samples for endosperm, diluting the proportion of pericarp sampled. We compared genotype results in an F2 population from B73 x Mo17 by first seed-chipping 96 F2 kernels and then germinating and leaf-sampling their corresponding, 96 F2 seedlings. DNA was prepared using the SDS method for both seed chip and leaf tissue and PCR was performed as above (Procedure, Section D). For 88 of the 96 individuals, sufficient PCR product was detected to call genotypes for both the chip- and leaf-based methods. Among the 88 individuals we focused on the homozygotes because in F2 seed-chip DNA, either homozygous genotype could theoretically be miscalled as heterozygous due to PCR amplification of DNA extracted from the heterozygous, maternal pericarp tissue. 47 individuals were homozygous, as called by the leaf-based method. Among their corresponding chip-DNA genotypes, 43 had the homozygous call,

### General Notes

1. Plastic and metal nail clipping pliers were tested (Figure 5); we recommend the metal pliers because they are durable, and the blade can be removed for sharpening or replacement.
2. Sufficient chipped seed tissue (15-30mg) should be collected for adequate DNA for one to several genotyping reactions. Within that constraint, feep the chipped tissue pieces as small as possible because larger particles are more challenging to grind in the DNA extraction step; during the chipping process, chip more endosperm to dilute and minimize the amount of pericarp sampled.
3. Seedlings from chipped seeds grew slower, perhaps due to reduced mobilization of nutrients from the reduced volume of aleurone and endosperm. Thus, we also recommend not chipping more endosperm than required for adequate DNA yield (see note 2). Moreover, chipped seeds are susceptible to mold infection, especially if germinated in paper towels for example. Thus, treating or disinfecting the seed before planting may be warranted.
4. Additional details on installing and using the plate mapping software as a Google sheet plug-in are available through the Vollbrecht lab website (https://faculty.sites.iastate.edu/vollbrec/tissue-sample-plate-mapper). When printing plate maps using the software, navigate to the “Plate labels for printing” tab and execute the print command with these settings: Print Current sheet; Paper size Letter; Landscape orientation; Scale Normal (100%); Margins Normal; Formatting Show gridlines and Show notes; Page order Over then down; Alignment Horizontal Center; Alignment Vertical Center; Headers and Footers selections according to your preferences.

**Figure 5.**
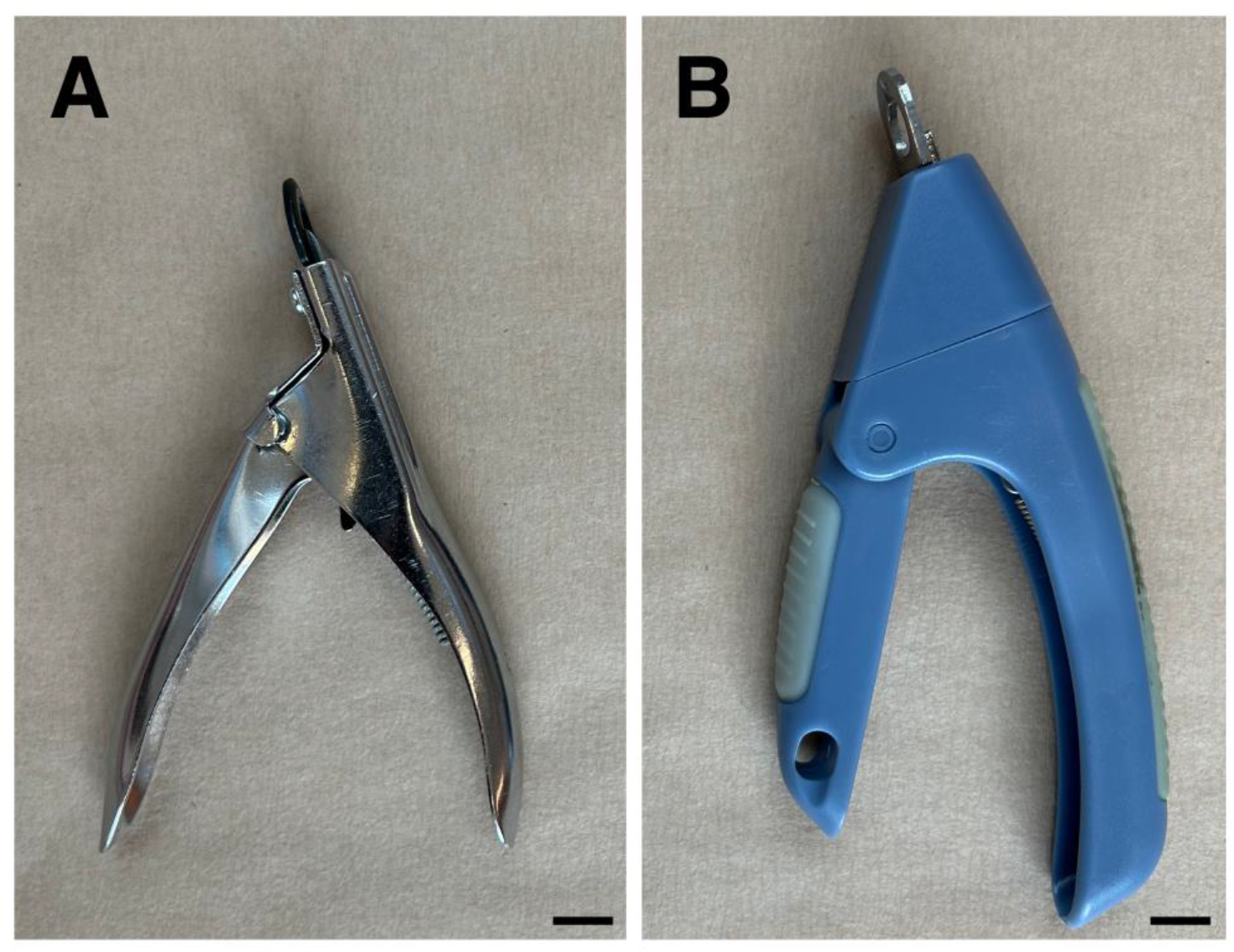
Nail clipping pliers tested. Clippers for seed chipping were either metal with exchangeable blade (A, preferred) or plastic with fixed blade (B).

## Supporting information

Supplementary protocol 1

Supplementary protocol 2

Supplementary protocol 3

## Acknowledgments

The tissue-sample plate mapper software was developed by Takao Shibamoto and Kokulapalan Wimalanathan with support from the National Science Foundation NSF-IOS grant 1238202 to E. Vollbrecht. Some figures were generated with BioRender.com. This research was supported by funding from the Iowa State University Crop Bioengineering Center.

## Competing interests

The authors declare no competing interests.

## Supplementary information

See Supplemental Protocol 1, Supplemental Protocol 2, and Supplemental Protocol 3.

## Notes

https://figshare.com/articles/media/Zebosi_et_al_seed_chipping_video_MOV/25529479

### Competing Interest Statement

The authors have declared no competing interest.

### Summary of Updates

Updated URL for supplementary movies; cleaned up protocols in Supplementary Information by stripping editing debris; manuscript content unchanged.

