## Supplementary protocol 1 for "An effective and safe maize seed chipping protocol using clipping pliers with applications in small-scale genotyping and marker-assisted breeding"

**Maize genomic DNA extraction using guanidine hydrochloride and filter plates**

Modified from (Gao et al., 2010)

### **Materials and Reagents**

**Biological materials**

1. Maize tissues (maize seeds and/or maize seedling leaf tissue)

**Reagents**

1. Sodium chloride, NaCl (CAS 7647-14-5)
2. Guanidine hydrochloride (CAS 50-01-1)
3. Tris-hydrochloride, Tris-HCl (CAS 1185-53-1)
4. Ethylenediaminetetraacetic Acid, EDTA (CAS 6381-92-6)
5. Ethanol (CAS 64-17-5)
6. HPLC grade Water (CAS 7732-18-5)

### **Recipes**

### 1. Wash buffer (See Recipes)

2. Extraction buffer (See Recipes)

### **Recipes**

1. Wash buffer
   a. 200 mM NaCl
   b. 50 mM Tris pH 7.4
   c. 70% ethanol
2. Extraction buffer
   a. 250 mM NaCl
   b. 4.2 M Guanidine-HCl
   c. 200 mM Tris-HCl pH 7.4
   d. 25 mM EDTA

**Laboratory Supplies**

1. AeraSeal breathable microplate sealing film (Excel Scientific, catalog number BS-25)
2. AcroPrep Advance 96-Well Filter Plate (Cytiva, catalog number 8032)
3. Strip Caps for 1.2 mL tubes (VWR, catalog number 82006-694)
4. 1.2 mL sampling individual tubes (VWR, catalog number 83009-678)
5. 2.2 mL 96-well sample collection plate (VWR, catalog number 43001-0020 or similar)
6. Parafilm
7. Pipette tips (200 µL and filtered 300 µL tips)
8. 0.8 mL 96 well plates (Thermo Scientific, catalog number AB-0859)
9. Stainless Steel Grinding Beads, 4 mm (Fisher, catalog number 2150)
10. Round bottom 96 well storage plate (Thermo Scientific, catalog number 12-565-502)
11. 96-well plate Microseal B Sealing Film (BioRad, catalog number MSB1001)

### **Equipment**

1. Tissue homogenizer (e.g. SPEX SamplePrep Geno/Grinder, Model 2010)
2. Benchtop centrifuge with swinging bucket rotor (e.g. Thermo Sorvall Legend XTR Centrifuge)
3. Multi-channel pipettes (100-300 μL) with extended tips

### **Procedure**

1. Collect up to 96 sampling tubes in one tissue block. Add to each sampling tube:
   a. 15-30 mg of chipped seed-endosperm or 2-3 leaf tissue punches, ~6 mm size
   b. 2 grinding beads
   c. 600 μL Extraction buffer for chipped seed (400 μL for leaf punches)
2. Seal sampling tubes well:
   a. place a layer of parafilm across the tube openings
   b. place strip caps on the parafilm and push them firmly into place
3. Grind tissue in Geno/Grinder at 1650 rpm (leaf tissue 1.5 minutes, endosperm 3 minutes).
4. Centrifuge at 3000 x *g* for 20 minutes at 10 ˚C.
5. Using filter tips, transfer 300 μL to the corresponding wells of a 0.8 mL plate. (**Critical**: avoid pipetting close to the pellet).
6. Seal the plate with Microseal film and centrifuge the 0.8 mL plate at 3000 x *g* for 10 minutes at 10 ˚C.
7. Using filter tips, transfer 250 μL from the 0.8 mL plate to the corresponding wells in the filter plate, avoiding any debris at the bottom of the 0.8 mL plate. (**Critical**: debris will clog the filter plate).
8. Seal the filter plate with an air permeable seal and place on a deep well 0.8 mL plate (same plate used in step 5) to collect the filtrate waste.
9. Centrifuge at 1500 x *g*, 5 minutes, at room temperature (RT) and discard the filtrate by pouring off.
10. Peel back the seal on the filter plate, add 200 μL Extraction buffer to each well, and re-seal it.
11. Centrifuge at 1500 x *g*, 5 minutes, RT, and discard the filtrate by pouring off.
12. Peel back the seal on the filter plate, add 200 μL Wash buffer to each well, and re-seal it.
13. Centrifuge at1500 x *g*, 5 minutes, RT, and discard the filtrate by pouring off.
14. Repeat steps 12 and 13.
15. Place the filter plate on an empty (waste) DNA storage plate, centrifuge at 1500 x *g*, 10 minutes, RT to remove residual wash. Discard the residual flowthrough.
16. To elute the DNA, place the filter plate on a new DNA storage plate. Peel back the seal on the filter plate, add 100 μL of HPLC grade water to each well, and re-seal it.
17. Centrifuge at 1500 x *g*, 5 minutes, RT, to elute the DNA into the storage plate. Seal the DNA storage plate with a Microseal film and store at 4˚C.
18. Discard the filter plate.
19. Use approximately 3 μL DNA in a 13 μL PCR reaction.
