## Supplementary protocol 2 for "An effective and safe maize seed chipping protocol using clipping pliers with applications in small-scale genotyping and marker-assisted breeding"

**Modified SDS-Based DNA Extraction Protocol**

Modified from (Edwards et al., 1991)

### **Materials and Reagents**

**Biological materials**

1. Maize tissues (maize seeds and/or maize seedling leaf tissue)

**Reagents**

2. Sodium chloride, NaCl (CAS 7647-14-5)
3. Tris-hydrochloride, Tris-HCl (CAS 1185-53-1)
4. Ethylenediaminetetraacetic Acid, EDTA (CAS 6381-92-6)
5. Sodium Dodecyl Sulfate, SDS (CAS 67-63-0)
6. Isopropanol (CAS 151-21-3)
7. Ethanol (CAS 64-17-5)
8. HPLC Water (CAS 7732-18-5)

### **Solutions**

### 1. SDS Extraction buffer (See Recipes)

2. 70% Ethanol

### **Recipes**

1. SDS Extraction buffer
   a. 200 mM Tris pH 7.5
   b. 250 mM NaCl
   c. 25 mM EDTA
   d. 0.5% SDS

**Laboratory Supplies**

1. Strip Caps for 1.2 mL tubes (VWR, catalog number 82006-694)
2. 1.2 mL sampling individual tubes (VWR, catalog number 83009-678)
3. 2.2 mL 96-well sample collection plate (VWR, catalog number 43001-0020 or similar)
4. Sealing Mat (Thermo Scientific, catalog number AB-0674)
5. Parafilm
6. Pipette tips (200 μL and filtered 300 μL tips)
7. Stainless Steel Grinding Beads, 4 mm (Fisher, catalog number 2150)
8. 0.8 mL 96 well plates (Thermo Scientific, catalog number AB-0859)
9. Round bottom 96 well storage plate (Thermo Scientific, catalog number 12-565-502)
10. 96-well plate Microseal B Sealing Film (BioRad, catalog number MSB1001)

### **Equipment**

1. Tissue homogenizer (e.g. SPEX SamplePrep Geno/Grinder, Model 2010)
2. Benchtop centrifuge with swinging bucket rotor (e.g. Thermo Sorvall Legend XTR Centrifuge)
3. Multi-channel pipettes (100-300 μl) with extended tips

### **Procedure**

1. Collect up to 96 sampling tubes in one tissue block. Add to each sampling tube:
   a. 15-30 mg of chipped seed-endosperm (or 1-2 leaf tissue punches, ~6 mm size)
   b. 2 metal grinding beads
   c. 600 μL of SDS Extraction buffer
2. Seal the sampling tubes well:
   a. place a layer of parafilm across the tube openings
   b. place strip caps on the parafilm and push them firmly into place
3. Grind tissue in Geno/Grinder at 1650 rpm (leaf tissue 1.5 minutes, endosperm 3 minutes).
4. Centrifuge at 3000 x *g* for 20 minutes at 10 ˚C.
5. During step 4, add 300 μL of isopropanol to wells of a 0.6 mL 96-well plate.
6. Using filter tips, transfer 400 μL of the tissue extraction supernatant (**Critical**: avoid the tissue pellet) into the isopropanol plate prepared in step 5.pipette up/down several times to mix.
7. Seal the plate with a Microseal film or a sealing mat and incubate at room temperature (RT) for 5 minutes.
8. Centrifuge at 3000 x *g* for 10 minutes at 10 ˚C.
9. Discard the supernatant by inverting the plate in one fluid motion. Keeping it inverted, gently tap dry on paper towels, then turn the plate upright and air-dry in the fume hood for 15 minutes.
10. Add 500 μL of 70% Ethanol to each well to rinse the DNA pellet (**Critical:** do not resuspend or disturb the pellet).
11. Seal the plate with a Microseal film or a sealing mat and centrifuge at 3000 x *g* for 10 minutes at 10 ˚C.
12. Discard the supernatant and air-dry the pellet plate in a fume hood for 30 minutes or until all ethanol has evaporated.
13. Add 120 μL HPLC water to each well of the pellet plate (**Critical:** do not resuspend or disturb the pellet).
14. Seal the plate with a Microseal film or a sealing mat and incubate for 10 minutes at RT to dissolve the DNA. With longer incubation time the pellet will release debris.
15. Centrifuge at 1500 x *g*, for 5 minutes, at RT.
16. Transfer 80 μL DNA solution into a storage plate while avoiding the pellet on the bottom. Seal the DNA storage plate with Microseal film and store at 4 ˚C.
17. Use approximately 3 μL DNA in a 13 μL PCR reaction.
