## Supplementary protocol 3 for "An effective and safe maize seed chipping protocol using clipping pliers with applications in small-scale genotyping and marker-assisted breeding"

**Urea-Based DNA Extraction Protocol**

Modified from (Leach et al., 2016).

### **Materials and Reagents**

**Biological materials**

1. Maize tissues (maize seeds and maize seedling leaf tissue)

**Reagents**

2. Sodium chloride, NaCl (CAS 7647-14-5)
3. N-Lauroyl sarcosine (CAS 137-16-6)
4. Glacial acetic acid (CAS 64-19-7)
5. Ammonium hydroxide (CAS 1336-21-6)
6. Tris-hydrochloride, Tris-HCl (CAS 1185-53-1)
7. Ethylenediaminetetraacetic Acid, EDTA (CAS 6381-92-6)
8. Sodium Dodecyl Sulfate, SDS (CAS 67-63-0)
9. Isopropanol (CAS 151-21-3)
10. Ethanol (CAS 64-17-5)
11. Double-distilled water
12. HPLC grade Water (CAS 7732-18-5)

### **Solutions**

1. 4.4 M NH_4_OAc, pH 5.2 (See Recipes)
2. Ammonium acetate : isopropanol solution, 1:10 (See Recipes)
3. Urea Extraction buffer (See Recipes)
4. 70% Ethanol

### **Recipes**

1. 4.4 M NH_4_OAc, pH 5.2 (to make 200 mL)
   a. 105 mL distilled water
   b. 50.5 mL Glacial acetic acid
   c. 45 mL 14.8N NH_4_OH
   d. Instructions: Working in a fume hood, add glacial acetic acid to the water and mix well. Then slowly add the NH_4_OH to the mixture, mixing the solution continuously by gently swirling the vessel while adding small aliquots of NH_4_OH.
2. Ammonium acetate : isopropanol solution, 1:10
   a. 450 ml isopropanol
   b. 50 ml 4.4 M ammonium acetate, pH 5.2
   c. Instructions: Working in a fume hood, add the ammonium acetate slowly to the isopropanol. Store at room temperature for up to 12 months.
3. Urea Extraction buffer
   a. 8M urea
   b. 50 mM Tris-HCl pH 8
   c. 0.35 M NaCl
   d. 20 mM EDTA
   e. 1 % n-lauroyl sarcosine

**Laboratory Supplies**

1. Strip Caps for 1.2 ml tubes (VWR, catalog number 82006-694)
2. 1.2 ml sampling individual tubes (VWR, catalog number 83009-678)
3. 2.2 mL 96-well sample collection plate (VWR, catalog number 43001-0020 or similar)
4. Sealing Mat (Thermo Scientific, catalog number AB0674)
5. Parafilm
6. Pipette tips (200 μL and filtered 300 μL tips)
7. 0.8 mL 96 well plates (Thermo Scientific, catalog number AB-0859)
8. Stainless Steel Grinding Beads, (Fisher, catalog number 2150)
9. Round bottom 96 well storage plate (Thermo Scientific, catalog number 12-565-502)
10. 96-well plate Microseal B Sealing Film (BioRad, catalog number MSB1001)

### **Equipment**

1. Tissue homogenizer (e.g. SPEX SamplePrep Geno/Grinder, Model 2010)
2. Benchtop centrifuge with a swinging bucket rotor (e.g. Thermo Sorvall Legend XTR Centrifuge)
3. Multi-channel pipettes (100-300 μL) with extended tips

### **Procedure**

1. Collect up to 96 sampling tubes in one tissue block. Add to each sampling tube
   a. 15-30 mg chipped seed-endosperm (or 1-2 leaf tissue punches, ~7 mm size)
   b. 2 grinding beads
   c. 600 μL Urea Extraction buffer
2. Seal the sampling tubes well
   a. place a layer of parafilm across the tube openings
   b. place strip caps on the parafilm and push them firmly into place
3. Grind tissue in Geno/Grinder at 1650 rpm (leaf tissue 1.5 minutes, endosperm 3 minutes).
4. Centrifuge at 3000 x *g*, 20 minutes at 10 ˚C.
5. During step 4, to each well of a 0.6mL plate add 300 μL of 1:10 NH_4_OAc:isopropanol (see recipes).
6. Using filter tips, transfer 400 μL of the tissue extraction supernatant (**Critical:** avoid the pellet) into the corresponding wells of the plate prepared in step 5 and pipette up/down several times to mix.
7. Seal the plate with a Microseal film or a sealing mat and incubate at room temperature (RT) for 5 minutes.
8. Centrifuge at 3000 x *g*, 10 minutes at 10 ˚C.
9. Discard the supernatant by inverting the plate in one fluid motion. Keeping it inverted, gently tap dry on paper towels, then turn the plate upright and air-dry in the fume hood for 15 minutes..
10. Add 500 μL of 70% Ethanol to each well to rinse the DNA pellet (**Critical:** Do not re-suspend or disturb the pellet).
11. Seal the plate with a Microseal film or a sealing mat and centrifuge at 3000 x *g* for 10 minutes at 10 ˚C.
12. Discard the supernatant and air-dry the pellet plate in a fume hood for 30 minutes or until all ethanol has evaporated.
13. Add 120 μL HPLC-grade water to each well of the pellet plate (**Critical:** Do not resuspend or disturb the pellet).
14. Seal the plate with a Microseal film or a sealing mat and incubate for 10 minutes RT to dissolve the DNA.
15. Centrifuge at 1500 x *g* for 5 minutes, at RT. .
16. Transfer 80 μL resuspended DNA into a storage plate while avoiding any pellet on the bottom. Seal the DNA storage plate with a microseal seal and store at 4 ˚C.
17. Use approximately 3 μL DNA in a 13 μL PCR reaction
